# Cell contractility-induced mechanical stress triggers eIF2α-regulated translation during early adhesion of mesenchymal-like cells

**DOI:** 10.1101/2022.04.19.485843

**Authors:** Alexia Caillier, Mathilde Lavigne, Jonathan Bergeman, Nadine Morin, Félix Rondeau, Juliette Gouhier, Benjamin Trudel, Jean-Philippe Lambert, Rachid Mazrouï, François Bordeleau, Marc-Étienne Huot

## Abstract

Cellular invasion is a complex process that requires several interdependent biological mechanisms, which are initiated by changes in adhesion that establish a morphology favorable for migration. Hence, the regulation of adhesion potential is a rate-limiting step in metastasis. Our previous work revealed that *de novo* translation is necessary to regulate the adhesion of mesenchymal-like cells; however, the underlying translational regulatory mechanism and the identity of newly synthesized proteins needed for the adhesion process remain unidentified. Here, we identify a mechanotransduction pathway linking force-mediated protein unfolding to translational reprogramming via the activation of the integrated stress response (ISR). Specifically, we demonstrate that phosphorylation of eukaryotic translation initiation factor 2 alpha (eIF2α) during early adhesion events leads to selective translation of mRNAs encoding proteins critical for adhesion complex formation, mechanosensing, and contractility. These findings uncover a mechanosensitive translational control axis that links intracellular force generation to cell adhesion and stress adaptation, with implications for understanding mesenchymal cell adhesion and morphological behavior

## INTRODUCTION

Cell adhesion is a fundamental biological process through which cells interact and attach to neighboring cells or the extracellular matrix (ECM) via specialized protein complexes. This process is mediated by a variety of adhesion molecules, including integrins, cadherins, selectins, and members of the immunoglobulin superfamily ^1^. These molecules not only anchor cells in place but also facilitate communication between the intracellular and extracellular environments ^2,3^. Cell adhesion plays a critical role in numerous physiological processes such as, tissue formation and maintenance during embryogenesis and wound healing ^4^, immune response ^5^, signal transduction ^6^, and cancer progression ^1^. In the latter case, regulation of cell adhesion is a crucial step to start cancer cell migration and invasion, as they depend on the dynamic disassembly and formation of nascent focal adhesions, that can progressively mature into stable adhesion sites, allowing the cell to anchor to the ECM ^7^.

Focal adhesion maturation depends on intracellular forces generated by actin treadmilling and actomyosin-mediated contractility ^3^. Several focal adhesion proteins undergo conformational changes in response to an imposed mechanical load, which in turn allow recruitment of additional focal adhesion proteins to reinforce or activate signaling pathways ^2^. For instance, the vinculin-interacting rod domains on Talin switch to an open conformation at around 5 pN^8^. A single Talin molecules can support over 50 pN ^8^ whereas the traction stress exerted on a focal adhesion can reach 0.8 nN/µm^2^ ^9^. Furthermore, when adhesion forms, the traction stress within a focal adhesion initially increases as it matures before going back down ^9,10^. For a cell that just adhered to a substrate, the high level of traction stresses occurring across multiple focal adhesions maturing in parallel may be sufficient to result in large-scale unfolding of adhesion-related proteins. For a highly contractile tumor cell ^11^, the mechanical stress-mediated protein unfolding could pose a challenge in maintaining the pool of focal adhesion proteins.

The accumulation of a large number of unfolded proteins can create aggregate that interfere with normal cellular functions, resulting in cellular stress that must be resolved ^12^. There are accumulating evidence showing that force-mediated protein unfolding is sufficient to trigger this cellular stress response ^13^. For instance, in muscle cells subjected to cyclic stretch, the cytolinker Filamin can undergo a force-mediated unfolding that is sufficient to trigger the recruitment of unfolded-protein response proteins and subsequent targeting for autophagic degradation^14^. One of the central regulator of the unfolded-protein stress response is the eukaryotic translation initiation factor 2 subunit alpha (eIF2α) that controls global inhibition of protein synthesis^15^. Upon phosphorylation, eIF2α triggers a widespread translation inhibition, allowing the selective translation of stress-responsive proteins^14^. Interestingly, cyclic stretch also induces phosphorylation of eIF2α in fibroblast and muscle cells^16–18^. Thus translational regulation constitutes a key mechanism by which cells adapt to the stress they encounter^15^. Interestingly we have shown that impairing translation, using various antibiotics known to specifically inhibit translation, greatly impaired mesenchymal cells capacity to adhere to different substrates, including plastic, extracellular matrix components, and cultured cell monolayers^19^. This effect was also observed in highly metastatic cancer cell lines that had gone through the epithelial-to-mesenchymal transition where global translation inhibition decreased adhesion capacity^19^. In this context, the requirement for translation regulation in those cells to properly attach further suggests a connection with the integrated stress response (ISR).

Herein, we describe a mechanism involving a stress-mediated eIF2α response during cell adhesion that is triggered by intracellular forces and involves translational reprogramming that leads to the translation of specific mRNAs during adhesion. Our investigation shows that mesenchymal-like cells, including highly metastatic cancer cells, adapt their translational activity by selectively translating mRNAs coding for key proteins involved in adhesion complex formation. We identify eIF2α as the main regulator of this adaptive response, which modulates the translation of adhesion-related proteins, as well as protein involved in mechanosensing and cellular contractility, thus highlighting a link between mechanical forces and translational control exclusive to mesenchymal like cells.

## RESULTS

### Phosphorylation of eIF2α in adhering cells

We first proceeded to measure the activation of the stress response pathway by assessing the phosphorylation of eIF2⍺ at different time points during adhesion in mesenchymal-and epithelial-like cells. For this, we used MRC-5 primary human fibroblasts as a representative mesenchymal like cell line and HeLa cells as a comparative epithelial-like line, as we previously did ^19^. We also tested MDA-MB-231 cell line, which expresses mesenchymal markers and the MDA-MB-468 cell line, which expresses epithelial markers. We observed a phosphorylation of eIF2⍺ during early stages of adhesion (0-60 minutes post seeding) in both MRC-5 and MDA-MB-231 (Fig. 1, left panels), which gradually declined as adhesion continued. Conversely, we observed no variations in eIF2α phosphorylation during adhesion in either HeLa or MDA-MB-468 cells (Fig. 1, right panels). These results suggest that there is a cell-adhesion mediated stress response only occurring in mesenchymal and mesenchymal-like cells.

**Fig. 1.**
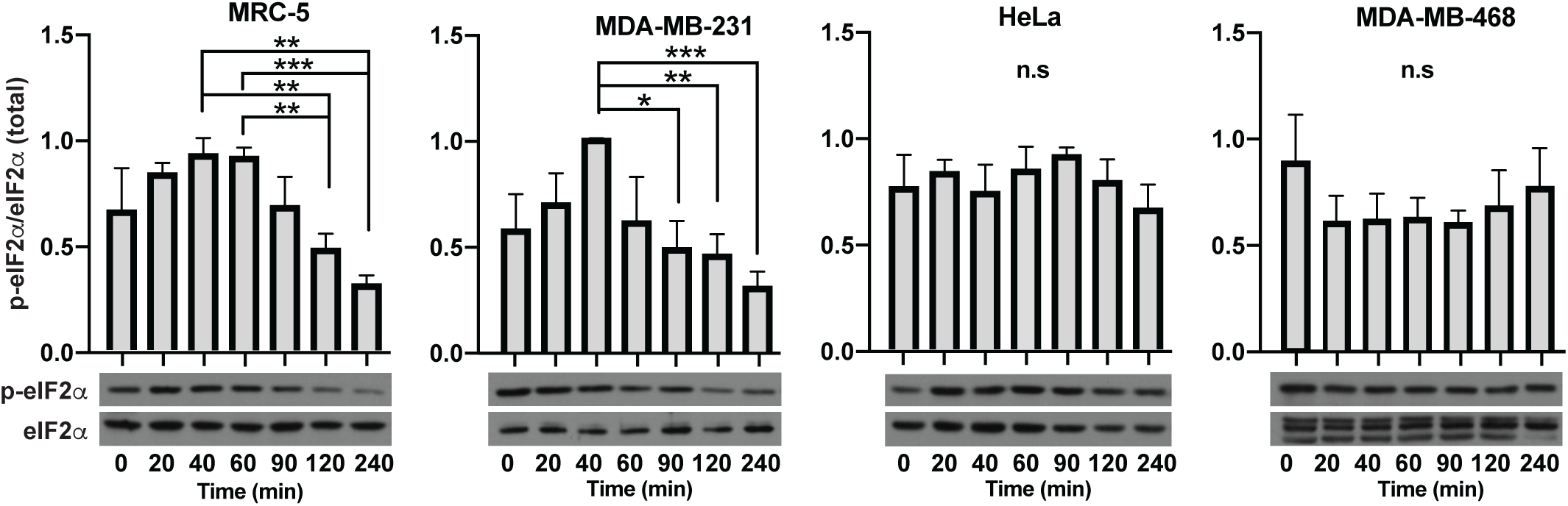
Phosphorylation of eIF2α during cellular adhesion. **(A**) Quantification of phosphorylated eIF2⍺ normalized to total eIF2α during the adhesion of MRC-5, MDA-MB-231, HeLa, and MDA-MB-468 cells. Assays were performed in triplicate. Error bars represent the standard deviation. n.s. *p* >0.05; \**p* ≤0.05; \*\**p* ≤0.01; \*\*\**p* ≤0.001; \*\*\*\**p* ≤0.0001 by two-tailed *t*-test.

### Translational dynamics during cell adhesion

Since eIF2α is the main initiator of translation ^20^, we then sought to assess if translation dynamics was altered in mesenchymal-and epithelial-like cells in response to this adhesion-mediated stress. In order to do so, we performed puromycylation assays, in which cells were treated with puromycin at different time points during adhesion to quantify their *de novo* translation activity ^21^. Indeed, during translation, puromycin is incorporated into nascent peptides and causes premature termination of protein synthesis and the release of the puromycilated polypeptides, thereby the amount of puromycilated polypeptides directly correlates with translational activity (Fig. 2B). As expected, we observed a significant decrease in translation activity in MRC-5 cells immediately after seeding, followed by an increase in translation as adhesion progressed (Fig. 2A). Conversely, we did not observe significant variations in translation throughout the HeLa adhesion process (Fig. 2B). The pattern of translational regulation in MRC-5 cells was confirmed by estimating active mRNA translation using polyribosomal profiling (Fig. S1). To confirm that the translation dynamics observed during the adherence process of MRC-5 cells were not cell-type specific, but rather dependent on cell morphology, we next examined two cancer cell lines known for their distinctive morphology and invasiveness ^22–24^. In concordance with the mesenchymal cell line (MRC-5), MDA-MB-231 cells displayed a translational repression during the initial adhesion process, followed by gradual translational activation (Fig. 2C). Like HeLa cells, we observed no variations in MDA-MB-468 cell translation during adhesion (Fig. 2D). Collectively, our results suggests that the adhesion-mediated translational regulation occurs only in mesenchymal-like cells and correlates with the associated stress.

**Fig. 2.**
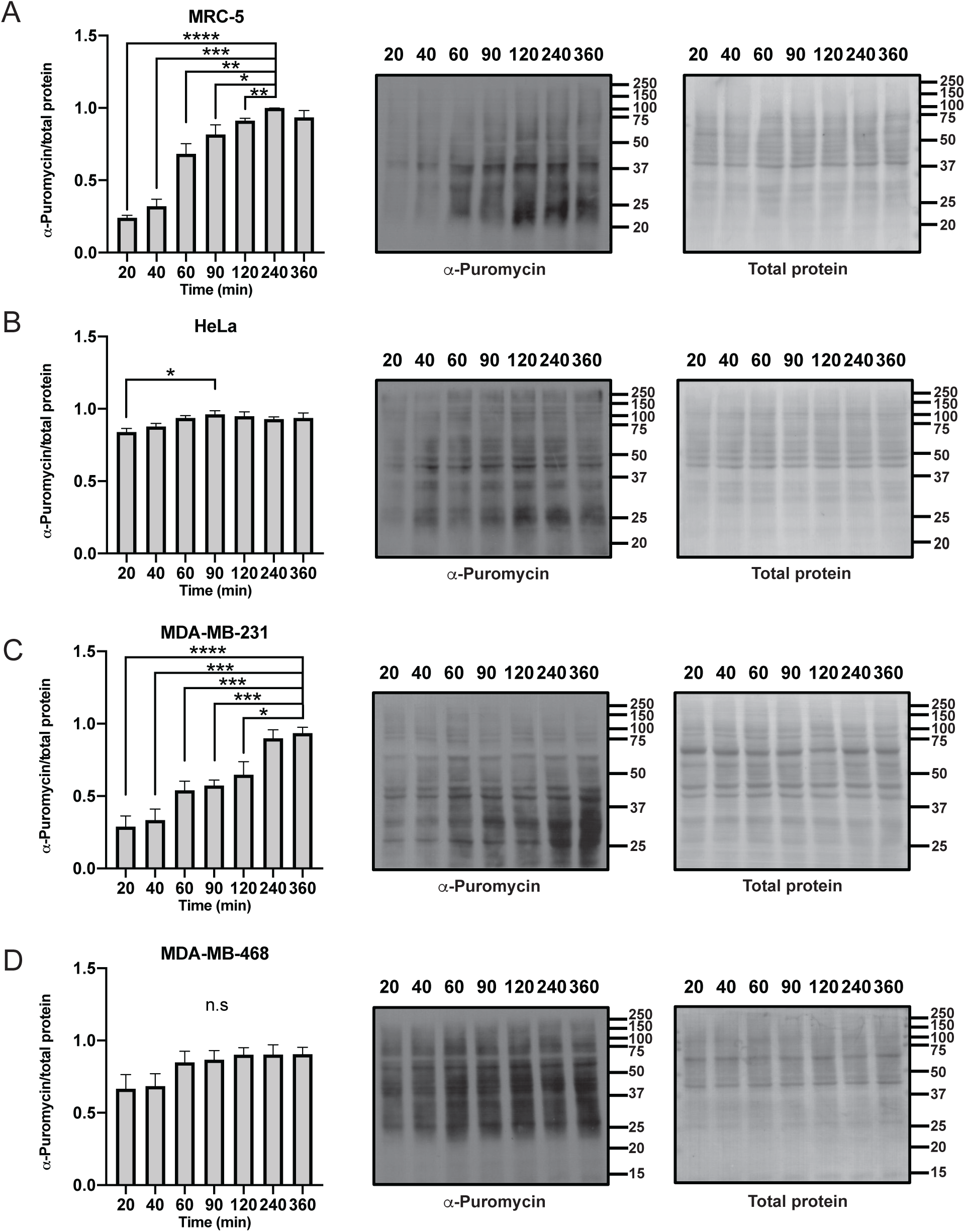
Translational dynamics during cellular adhesion Puromycin incorporation within actively synthetized polypeptides (5 mg/mL puromycin / 5 min) in (**A**) MRC-5, (**B**) HeLa, (**C**) MDA-MB-231, and (**D**) MDA-MB-468 cells. Puromycilation was revealed using an anti-puromycin specific antibody (12D10). Graphical representations (left panels) show the puromycin signals (middle panels) over the normalized amido black signals (right panels). Assays were performed in triplicate. Error bars represent the standard deviation. n.s *p* >0.05; \**p* ≤0.05; ** *p* ≤0.01; *** *p* ≤0.001; \*\*\*\**p* ≤0.0001 by two-tailed *t*-test.

### Phosphorylation of eIF2α modulates adhesion-related translation

To fully establish if this translation retrogradation during mesenchymal-like cells adhesion is directly linked to eIF2α stress response, we treated cells with 1 µM ISRIB (Integrated Stress Response Inhibitor), to bypass eIF2α phosphorylation-mediated translational inhibition. When eIF2α is phosphorylated, it acts as a competitive inhibitor of eIF2B, blocking its guanine nucleotide exchange activity, which is required for translation initiation. The synthetic bisglycolamide ISRIB directly binds eIF2B, rendering its activity independent of p-eIF2α control and restoring translation under stress^25,26^.

As shown by puromycin incorporation assays (Fig. 3A), ISRIB treatment abrogated the translational inhibition normally observed in these cells (Fig 2A and D). Moreover, we observed increased relative adhesion over time of both MRC-5 and MDA-MB-231 cells following ISRIB pretreatment. In contrast, cycloheximide-induced translational repression decreased the adhesion kinetics in both cell lines (Fig 3B). These results further support that the adhesion related translational pattern observed in mesenchymal cell is controlled through an eIF2α stress response.

**Fig. 3.**
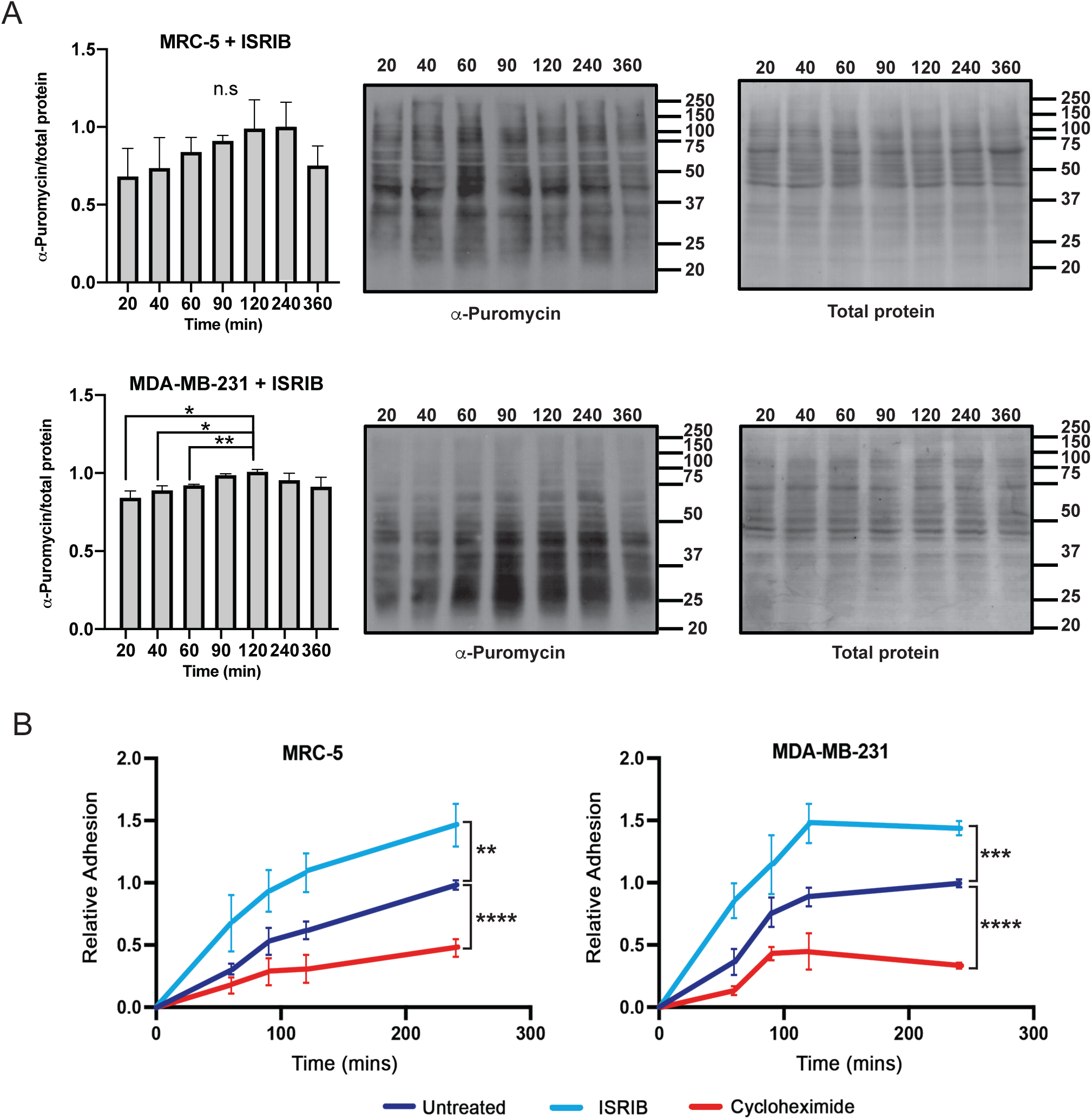
Phosphorylation of eIF2α regulates adhesion-related translation. (A) Puromycilation assays (5 mg/mL puromycin / 5 min) on MRC-5 cells pretreated with 1 μM ISRIB (60 min). Graphical representations (left panels) show the puromycin signals (middle panels) over the normalized amido black signals (right panels). Assays were performed in triplicate. Error bars represent the standard deviation. n.s p >0.05; *, p ≤0.05; **, p ≤0.01; ***, p ≤0.001; ****, p ≤0.0001 by two-tailed t-test. (B) In vitro adhesion assays with MRC-5 (n=3) and MDA-MB-231 (n=2) cells treated with ISRIB (1 μM) or cycloheximide (50 mg/mL) or untreated (DMSO).

### Identification of the adhesion-regulated proteome

Considering this stress-related regulation of translation, we next proceeded to determine which proteins were regulated. To investigate this, we identified proteins specifically translated 45 minutes post-seeding, at which point the largest increase in translation during adhesion was observed. To identify newly synthetized proteins, we immunoprecipitated puromycinated peptides (Fig. 4A and B). MRC-5 cells were allowed to adhere to the substrate for 40 min, which correspond to the time point where eIF2α phosphorylation start to decrease (Fig. 1), and then treated with puromycin for an additional 5 min. 48 h post-seeding MRC-5 cells were treated with puromycin for 5 min as the baseline. Puromycilated peptides were immunoprecipitated from both samples (Fig. 4A) and identified by mass spectrometry. Proteins were considered preferentially synthetized in adhering cells if they had three or more peptides with scores at least ten-fold higher than in the control in at least two biological triplicates. Interestingly, when we investigated the cellular components or molecular functions that were enriched within the newly synthesized proteins, then main GO terms that came out were associated with the focal adhesion, cell-cell junctions and the cytoskeleton assembly (Fig 4C and D). Unsurprisingly, GO terms associated with the ribosome were significantly enriched (Fig 4C and D). Of note, GO terms associated with the unfolded protein response also came out as significant (Fig 4C). Specifically, we identified 47 proteins that were specifically synthesized in adhering cells, among which 33% are already known to be directly involved in focal complex formation, maturation, and anchoring to the forming actin cytoskeleton (Fig. 4E). These results suggest that the translation pattern observed in mesenchymal-like cells during adhesion results in the redirection of the translational machinery toward transcripts encoding adhesion-related proteins.

**Fig. 4.**
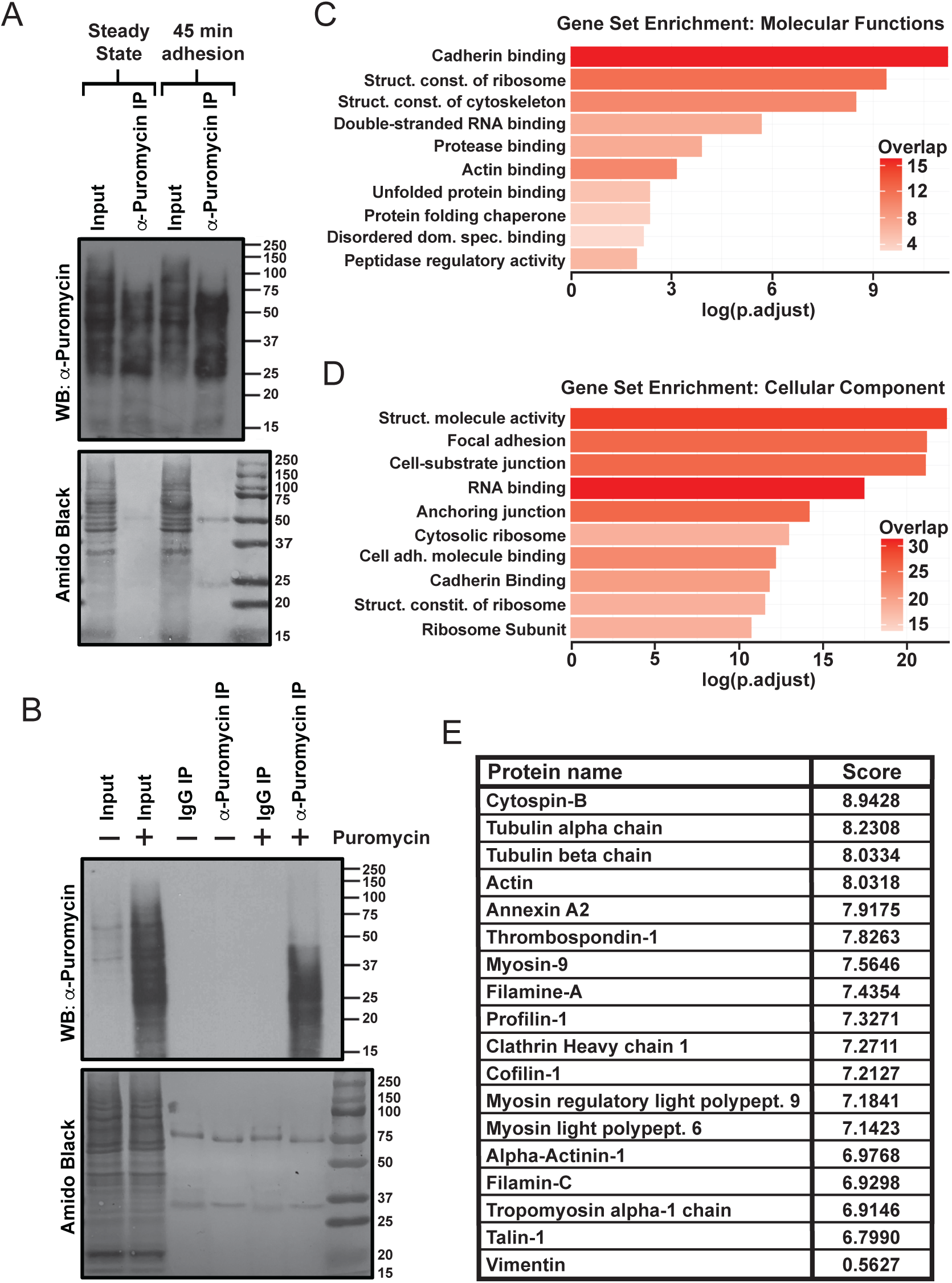
Identification of the adhesion-related proteome (**A**) Representative a-puromycin (clone 12D10) immunoprecipitation of puromycin-treated steady state and adhering cells (45 min post-seeding). The extracts shown are representative of the triplicate immunoprecipitation of puromycin treated cells at steady state (fully adhered) and adhering cells (45 minutes post-seeding) used for peptide identification. (**B**) Puromycin immunoprecipitation on extract made from cells treated with either DMSO or 5 mg/mL puromycin for 5 minutes. Immunoprecipitation was also performed on DMSO treated cells (-Puromycin), as a negative control. Puromycilated polypeptides were immunoprecipitated using the anti-puromycin (clone 12D10) or control IgG antibodies and detected using anti-puromycin antibody. (**C**) Functional enrichment analysis associated with molecular functions and (**D**) cellular components GO terms of the translated proteins. Selection was based on the level of puromycilated peptides (10X higher) present in each sample, which correspond to actively translated mRNAs. Data is presented as adjusted p-value and color coded according to the number of detected proteins overlapping with the respective GO terms. (**E**) Summary table listing proteins involved in focal complex formation/maturation, anchoring to the forming actin cytoskeleton and the establishment of cellular morphology that were identified as being translated 45 minutes post-seeding identified by MS/MS following puromycin immunoprecipitation.

### Translational dynamics of adhesion-related proteins in mesenchymal cells

We then proceeded to validate the puromycin immunoprecipitation results and assess whether the specific translation of mRNAs encoding for adhesion proteins was unique to mesenchymal-like cells.

This was performed in adhering MDA-MB-231 by performing western blot and comparing protein levels over the course of the adhesion process. We selected specifically synthesized key proteins with involvement in the adhesion process (Talin) (Fig 5A), as well as proteins involves in the actin anchorage and contractility (Filamin and ⍺-Actinin) (Fig 5 B and C). In most cases, adhesion proteins were barely detectable in unadhered cells, and their expression increased gradually as adhesion was initiated (20–60 min) and consolidated (60–240 min). To determine if the gradual expression pattern of adhesion proteins was regulated by eIF2α phosphorylation, we forced translation by pretreating MDA-MB-231 cells with 1 µM ISRIB, thus bypassing eIF2α phosphorylation-mediated translational inhibition. ISRIB treatment completely disrupted the translation patterns of these proteins in MDA-MB-231 cells (Fig. 5). Finally, no changes in the expression of Talin, Filamin or ⍺-Actinine were observed in epithelial-like MDA-MB-468 cells (Fig S2), whereas differential expression of these proteins was detected in mesenchymal MRC-5 cells, as seen with the MDA-MB-231 (Fig S3). These results suggest that this translational regulatory mechanism is involved in the adhesion process of mesenchymal-like cells, but not in epithelial-like cells.

**Fig. 5.**
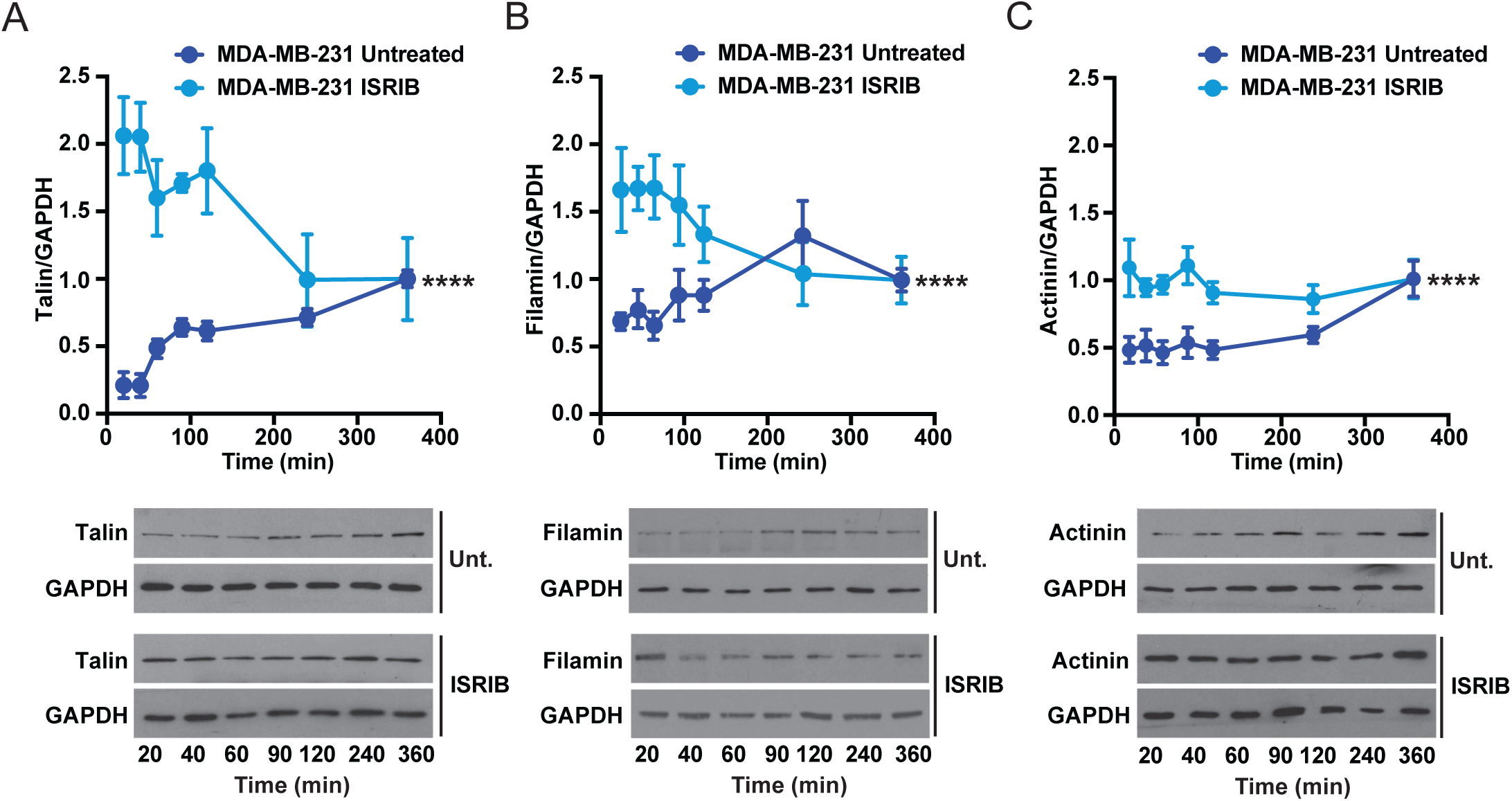
Expression of key adhesion-and cytoskeleton-related proteins during cell adhesion Quantification of levels of translated adhesion related mRNAs of Talin (**A**), Filamin-A (**B**) and α-Actinin (**C**) at different time points during cellular adhesion in untreated MDA-MB-231 cells (DMSO) or treated with 1μM ISRIB (n=3). Lower panels show representative western blot detection of each protein at different time points during cellular adhesion in untreated MDA-MB-231 cells (DMSO) and in MDA-MB-231 cells with 1µM ISRIB. Protein expression was normalized at time point 360 minutes, as we found that adhered cells were morphologically mature independently of treatments. Error bars represent the standard deviation. n.s *p* >0.05; \**p* ≤0.05; ** *p* ≤0.01; *** *p* ≤0.001; \*\*\*\**p* ≤0.0001 by two-way ANOVA

### Impact of translation on cellular contractility and traction forces during adhesion

Cell-generated forces are critical for the assembly and maturation of focal adhesions (FAs) ^3^. In turn, mechanical forces can induce a stress response that could repress translation, notably by inducing a force-drive protein unfolding ^27^. In this context, we investigated whether cell-generated force during cell adhesion was sufficient to trigger protein damage in line with the observed stress response. To do so, we started by implementing a computational approach based on the tensegrity model. The biophysical parameters that were included within the model were based on cell spreading rate, actomyosin generated tension, the stiffness of cytoskeletal bundles and cytoskeleton connectivity evolution over time (nodes in contact with the surface can be approximated to an adhesion) (Table S1, Fig. 6A-C). The model was then used to compute the force in each individual cytoskeletal components which, once integrated over the entire structure, highlighted a gradual increase of total force that appears to converge toward a plateau (Fig. 6D). Interestingly, when we looked at either the normal stress σ or shear stress τ_xy_, the model predicted that the strain behaved as an under-damped system, with a high peak that then dropped sharply before coming back up and overshooting the final equilibrium value (Fig. 6E and F). Of note, τ_xy_ was higher on average than σ. Next, we used the data on the shear stress and modeled number of adhesions as a function of time in a modified linear clutch model of the focal adhesion^28^ to determine the impact of the peak strain on a virtual pool of 30,000 Talin molecules. A peak threshold force of 100 pN applied to a single Talin was considered as sufficient to results in an unfolded state that may trigger an unfolded protein response and subsequent degradation. In addition, a maximal translation rate of 3000 Talin min^−1^ modulated by the p-eIF2α signal was assumed (∼60 to 160 proteins·min^−1^ produced per mRNA; typical 30 mRNA copies per gene) ^29,30^. Under these conditions, we could observe a decrease in the number of available Talin molecules over the first 20 iterations of the simulation, reaching a minimum that went back up once the stress per focal adhesion was low enough to avoid extreme traction stresses (Fig. 6G). Overall, these results suggest that the traction stresses that build up during cell adhesion are sufficient to deplete load-bearing proteins.

**Fig. 6.**
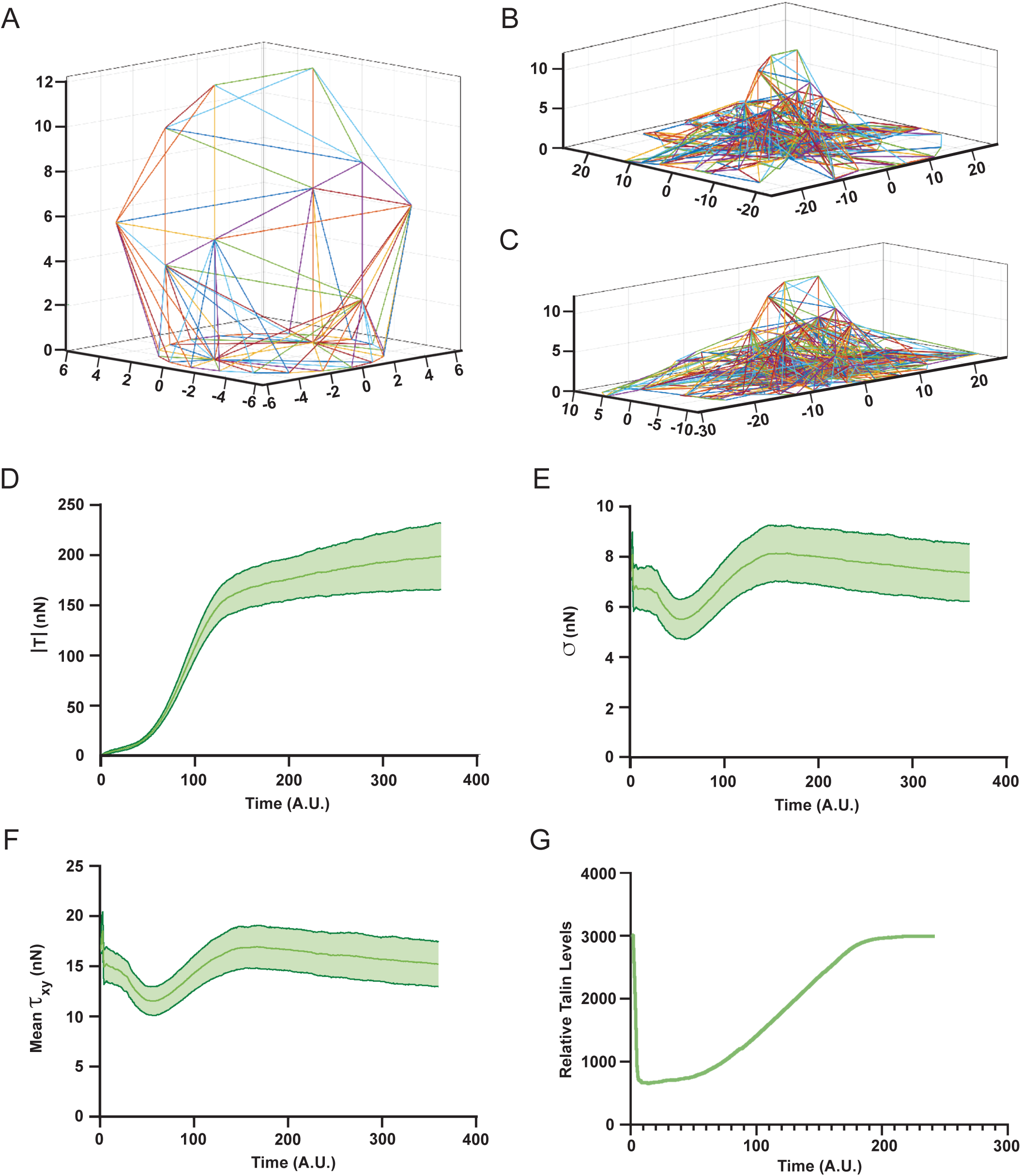
Computational modelization of cellular contractility and traction forces during mesenchymal-like cell adhesion. Computed organization of the tensegrity scaffold within a representative cell spreading on the substrate at (**A**) T = 0 min, (**B**) 60 min and (**C**) 360 min. (**D**) Average traction force |T| exerted by the spreading cell computed over 60 independent simulations along with the computed (**E**) normal (σ) and (**F**) shear (τ) stress within the elements of the scaffold. (**G**) Corresponding cellular content for Talin predicted by the clutch model assuming a failure threshold of 100 pN for a single Talin molecules. Error bands correspond to SD from 60 independent simulation.

We therefore sought to validate our model prediction and assess the impact of translational regulation on various mechanobiological parameters, including intracellular contractility and traction force generation. To assess intracellular contractility, we used quantitative polarization microscopy (QPOL) which provide a readout of cell contractility based on the impact of mechanical strain on the optical property of the sample ^31,32^. In MDA-MB-231 cells, we observed a decrease in the retardance signal indicative of a lower actin tension within the cell during the early phases of adhesion, followed by an increase of internal tension that peaked at 240 min before coming back down (Fig. 7A and B). Interestingly treatments with either ISRIB or cycloheximide prevented most of the temporal fluctuation of the retardance signal (Fig. 7B and Fig. S4). Next, we assessed traction force generation using real time traction forces microscopy (TFM). Similar to what was predicted by the model, the force generated by the cells was found to gradually increase over the course of the experiment and seemed to converge toward a plateau (Fig. 7C and D). In this case however, treatment with ISRIB resulted in a rapid increase of the measured traction forces at early time points (t < 60 min) followed by a slow decrease over time (Fig. 7D). In the case of cycloheximide, the treatment resulted in an apparent measured traction force that was consistently lower than the untreated control (Fig. 7D).

**Fig. 7.**
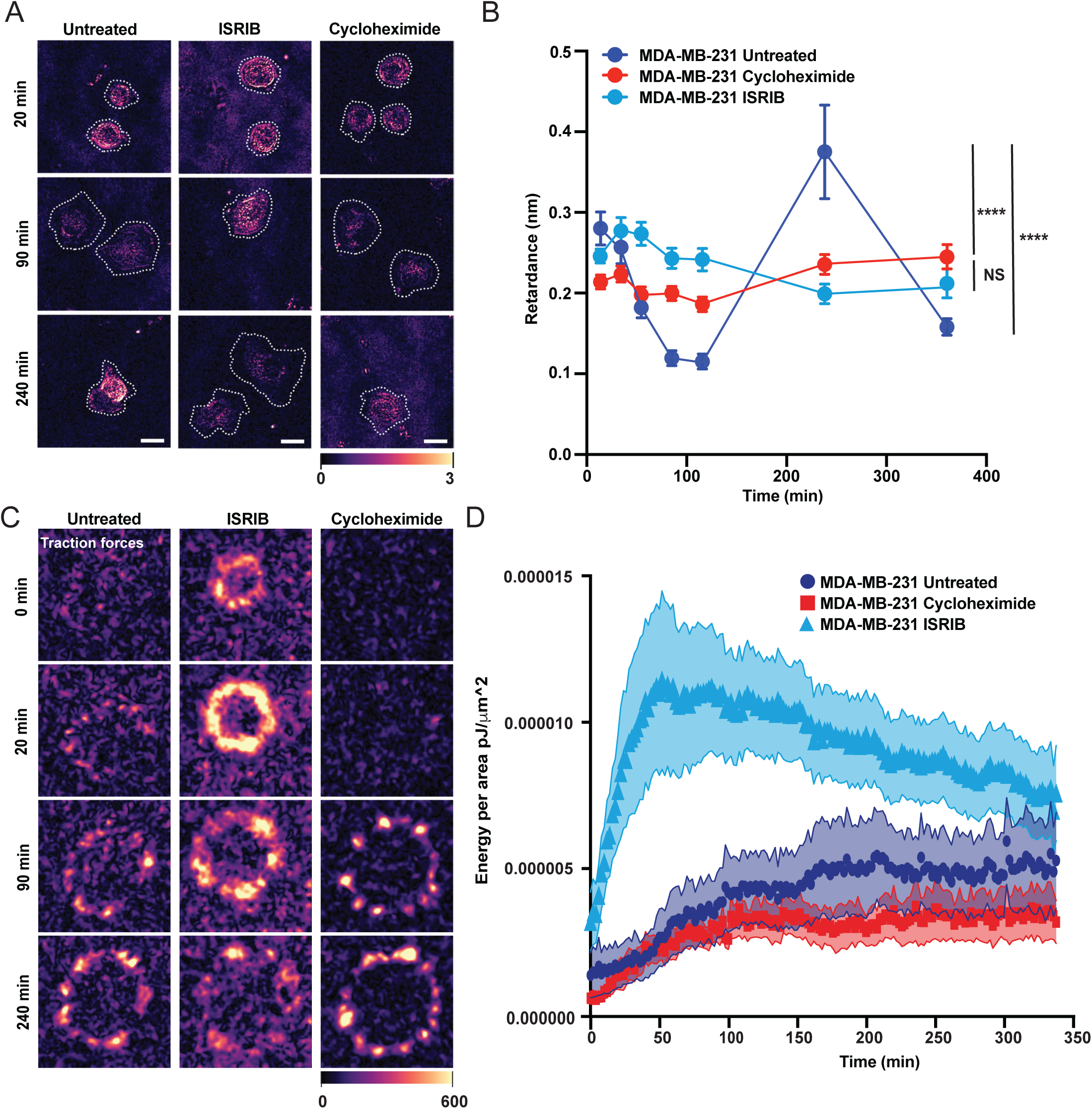
Impact of translation on cellular contractility and traction forces during adhesion (**A**) Representatives retardance images of untreated MDA-MB-231 cells (DMSO) or MDA-MB-231 treated with either cycloheximide (50μg/mL) or ISRIB (1μM) at 20 min, 90 min and 240 min post-seeding. Scale bar = 20 mm. (**B**) Measurements of retardance in untreated MDA-MB-231 cells (DMSO) or MDA-MB-231 treated with cycloheximide (50μg/mL) or ISRIB (1μM) after adhesion. (untreated 20min n=136, 40min n=97, 60min n=110, 90min n=108, 120min n=82, 240min n=115, 360min n=107; ISRIB 20min n=115, 40min n=86, 60min n=85, 90min n=81, 120min n=68, 240min n=56, 360min n=30; Cycloheximide 20min n=94, 40min n=106, 60min n=74, 90min n=89, 120min n=79, 240min n=81, 360min n=78). (**C)** Representative image of traction forces microscopy of untreated MDA-MB-231 cells (DMSO) or MDA-MB-231 treated with cycloheximide (50μg/mL) or ISRIB (1μM). (**D**) Real-time quantification of cellular traction the analyzed image of traction forces microscopy of untreated MDA-MB-231 cells (n=2, 11 cells total) or MDA-MB-231 treated with cycloheximide (n=2, 10 cells total) or ISRIB (n=3, 16 cells total). Data presented as mean ± SEM; error bars represent the standard deviation. n.s *p* >0.05; \**p* ≤0.05; ** *p* ≤0.01; \*\*\**p* ≤0.001; \*\*\*\**p* ≤0.0001 by two-way ANOVA Scale bar 10uM

Since both the computational model and experimental data indicated that the cytoskeleton and focal adhesions were initially under a tensional state, we proceeded to directly test if the mechanical stress was directly responsible for eIF2α activation. To do so, we pretreated the cells with Y27632, a selective inhibitor of Rho-associated protein kinases upstream of actomyosin contractility, to lower the tension within the cytoskeleton ^33^. As expected, Y27632 caused a progressive decrease of the optical retardance over time that was associated with an increased cell spreading compared to untreated cells (Fig. 8 A and B). The peak of internal tensional stress visible at 240 minutes post-adhesion in the controls was also abrogated following Y27632 treatment (Fig. 8A and B). We next compared the kinetics of eIF2α phosphorylation during the adhesion process, in presence or absence of Y27632 to confirm that internal tension was triggering the stress-induced response. As previously shown, the initial decrease in internal tension increased eIF2α phosphorylation (20-40 min post-seeding) in untreated cells, which was then followed by a slow decrease of the stress response (Fig. 8C and D). On the other hand, inhibition of contractility directly prevented the initial post-adhesion increase of eIF2α phosphorylation and only the decay over time was observed (Fig. 8C and D). Altogether, our results demonstrate that the increase in internal tension that follows initial adhesion induces a rapid stress response, associated with a loss of focal adhesion and actin-associated proteins, which then force adhesion proteins expression to sustain a mesenchymal morphology.

**Fig. 8.**
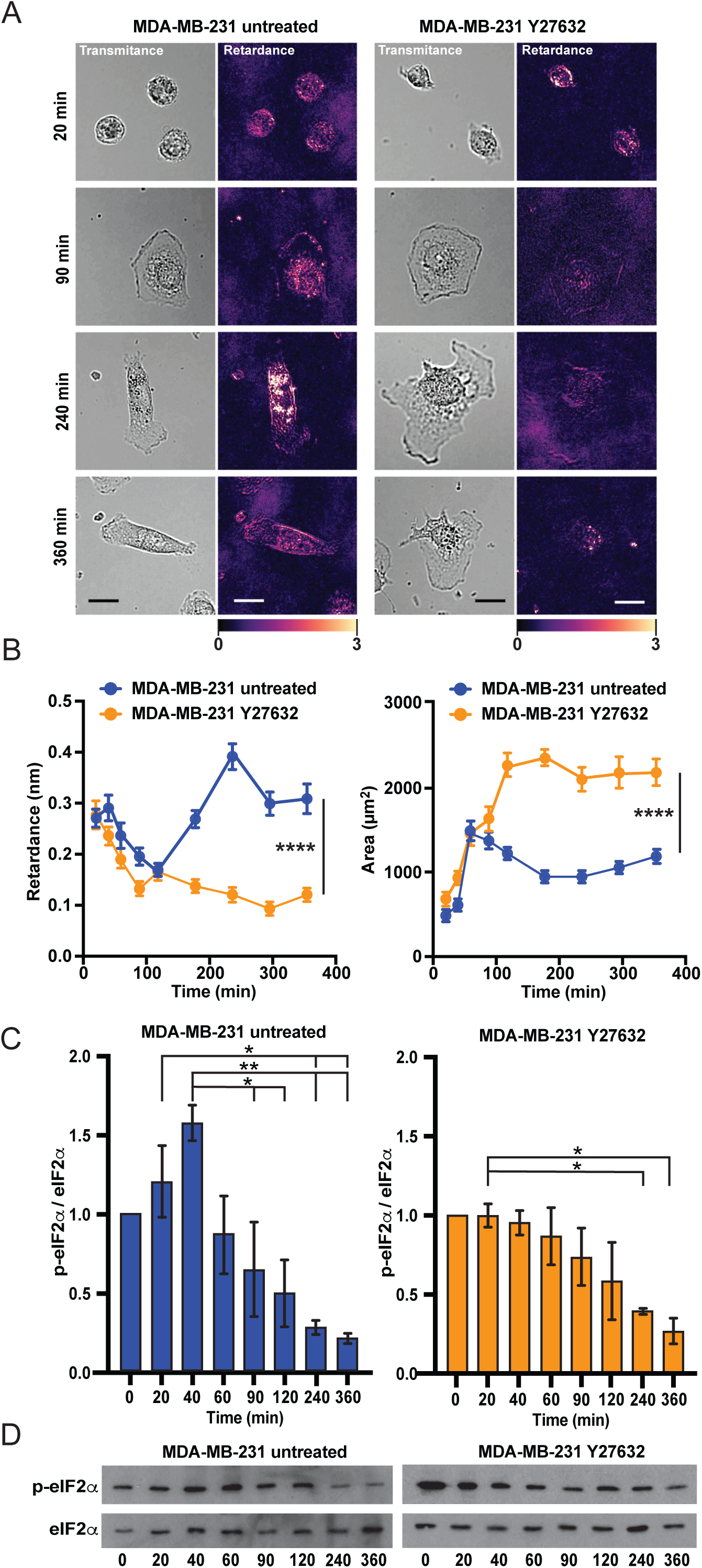
Mechanical stress induces eIF2α phosphorylation through cellular contractility. (**A**) Representative transmitted light (Transmitance) and retardance (Retardance) images of MDA-MB-231 cells untreated (left panel) or treated with Y27632 (right panel). (**B**)Measurements of retardance and cell area in untreated MDA-MB-231 cells (DMSO) and Y27632-treated with MDA-MB-231 cells (10μM) following seeding (untreated 20min n=83, 40min n=67, 60min n=69, 90min n=71, 120min n=77, 180min n=77, 240min n=64, 300min n=73, 360min n=64; Y27632 20min n=48, 40min n=75, 60min n=59, 90min n=52, 120min n=71, 180min n=67, 240min n=55, 300min n=42, 360min n=49). (**C**) Quantification of phosphorylated eIF2α normalized (p-eIF2α) on total eIF2α during the adhesion of untreated MDA-MB-231 cells (DMSO) or following treatment with 10μM Y27632 (n=3). (**D**) Representative detection of phosphorylated eIF2α (p-eIF2α) and eIF2α of MDA-MB-231 of untreated. Data presented as mean ± SEM; error bars represent the standard deviation. n.s *p* >0.05; \**p* ≤0.05; ** *p* ≤0.01; \*\*\**p* ≤0.001; \*\*\*\**p* ≤0.0001 by two-way ANOVA.

## DISCUSSION

Cell adhesion is a tightly regulated process governed by both extracellular and intracellular signals. These cues orchestrate the recruitment and activation of key proteins involved in the assembly of adhesion complexes, which, under favorable conditions, can mature into focal adhesion sites^3^. In this study, we demonstrated that an integrated stress response is activated following cell adhesion in mesenchymal-like cells followed by a modulation of general translation. Notably, cell-mediated contractility could trigger eIF2⍺ phosphorylation and thus repress translation. Moreover, this stress response was linked to subsequent preferential translational regulation of Talin, as well as other adhesion and structural proteins (e.g., α-Actinin and Filamin), notably through the action of intracellular mechanical strain. We found that this response was absent in epithelial cells, but present in mesenchymal-like cells. Finally, close control of the stress-response machinery in turn influenced cell-generated forces, implying that protein homeostasis and unfolded protein response is required under physiological conditions to maintain the contractile state of mesenchymal-like cells.

Interestingly, several of the strongest hits that were identified amongst the newly synthesised protein screen following cell adhesion are proteins that are deemed mechanosensitive. Importantly, the conformational changes that occurs upon the application of mechanical force can expose hidden interaction domains and modulate the functional roles of mechanosensitive proteins^2^, similar to how Talin recruits Vinculin^34^. For instance, Filamin-A and Filamin-C are known to unfold under the action of mechanical stress^35,36^, and this stretching is important for the modulation of YAP mechanotransduction^18^. Moreover, force-mediated Filamin-A unfolding can directly results in its degradation^37^. The α-Actinin homodimers can also stretch under load, and heterodimers can even break for load above 60 pN^38^. Myosins themselves are mechanosentive proteins^39,40^, that are able to change conformation when under mechanical load^41^. Even the Clathrins that are involved in intracellular trafficking and endocytosis undergo unfolding changes resulting from the mechanical load applied through Actin^42^. Thus, the unfolding of proteins acts as a source of stress, potentially interfering with their structural integrity and functional roles, and activating stress-related signaling cascades. Moreover, our results highlight that this can in fact involve large scale unfolding events in a large variety of these mechanosensitive proteins which triggers the ISR. It is likely that such a mechanism plays a critical role in regulating cell behavior beyond adhesion. Indeed, studies in mice have shown that suppression or partial disruption of the ISR was sufficient to hinder the progression of lung adenocarcinoma^43^.

Our computational model prediction and experimental results shows that initial adhesion increase internal tension, likely resulting in the unfolding of Talin and the other cytolinker Filamin and ⍺-Actinin. This then create a second cellular stress, as shown by the increased eIF2α phosphorylation during early adhesion, which can be abolished by inhibiting cellular contractility. While this directly implicated cellular contractility as a driver of stress occurring during cell adhesion, it remains that this is observed in mesenchymal-like cells only. The eIF2α-regulated response was not observed cancer cell lines that retained epithelial morphology marker expression. This could be in part due to difference in the organization and composition of the cytoskeleton as well as how cell contractility pathways are regulated. For instance, mechanical load is preferentially applied to cortical actin in epithelial cells whereas the load bearing elements in mesenchymal cells are the actin stress fibers^44,45^. As such, the orientation of the load-bearing elements relative to focal adhesions or actin architecture could influence the actual strain they are under^46^. Moreover, expression of the intermediate filament Vimentin is predicted to alter intracellular tensional stress ^47^. In addition, decreased expression of keratins (i.e. the epithelial intermediate filaments) reduces a cell’s ability to fine tune its contractility^48^. However, we cannot exclude the possibility that more direct changes in cell contractility associated with a mesenchymal phenotype could contribute to the observed difference in the stress response.

Following stress, cells can rapidly shutdown general translation initiation through a mechanism involving the phosphorylation of eIF2α ^49,50^ and concomitantly allows the translation of selected mRNAs encoding cell survival proteins ^51–53^. In fact, the main mRNAs that can be translated under stress conditions are heat shock proteins and other related chaperones ^14^. These likely precede the expression of the mechanosensitive proteins that we observe after the resolution of eIF2α translation repression and could well be related to the cell’s efforts to repair damaged proteins. Interestingly, several of the mechanosensing protein that are impacted during cell adhesion, including Filamins, α-Actinin and Myosin, rely on the chaperone-assisted selective autophagy to maintain their homeostasis^54–56^. The induction of the selective autophagy machinery that includes BAG3 and HSP70 by a proteotoxic stress can indeed induce eIF2α phosphorylation ^57^. This machinery is also reliant on actin dynamics while being involved in the resolution of the mechanical stress ^37,54^. Interestingly, the mechanosensitivity of MEFs and A7r5 cells following activation of RhoA appears to involve the chaperone-assisted selective autophagy system ^37^. Therefore, it is plausible that chaperone-assisted selective autophagy might play a critical role in detecting and resolving the tensional-related stress that occurs following cell adhesion we observed here. In line with this, we did detect unfolded proteins response chaperone, something that will warrant further investigation. In any case, resolving this stress response seems to be a clutch moment in cell adhesion that allows the translation of adhesion related mRNAs essential for adhesion consolidation and cell spreading, mechanosensing as well as proteins important for the morphological changes associated with cellular invasion ^11^.

Overall, these results clearly show that cytoskeletal stress induce eIF2α phosphorylation and dependent translational regulation during early adhesion, and that this mechanism is involved in mitigating force-induced protein depletion. Notably, our findings suggest that translational regulation during cell adhesion must be tightly controlled in cells adopting a mesenchymal morphology to compensate for internal tension stresses. Considering the importance of adhesion turnover in several physiological processes, most notably during metastasis, mechanical stress will be significant in mesenchymal-like tumor cells during cell migration, invasion or transmigration from the vascular system. The ability of cells that underwent EMT to handle the associated tensional stress could constitute a novel actionable target.

## AUTHOR CONTRIBUTIONS

Conceptualization: M.-E.H., F.B., A.C., J.B., and M.L.; Methodology: M.-E.H., F.B., A.C., M.L., N.M., B.T., R.M. and J.-P.L.; Validation: A.C. M.L. and N. M.; Formal analysis: M.-E.H., F.B., A.C., M.L., N.M., B.T., and J.-P.L..; Investigation: M.-E.H., F.B., A.C., M.L., N.M., B.T., R.M. and J.-P.L.; Resources: M.-E.H. and F.B., R.M. and J.-P.L.; Data curation: F.B., A.C., M.L., B.T. and J.-P.L.; Writing–original draft: M.-E.H., F.B., A.C., and M.L.; Supervision: M.-E. H. and F.B.; Project administration: M.-E. H. and F.B..

## DATA AVAILABILITY

Data will be made available on request.

## AKNOWLEDGEMENTS

We thank Carl St-Pierre and the Core Imaging Facility of the CHU de Québec–Université Laval Research Center (HEJ site) for their assistance with microscopic analyses. We thank Patrick Oakes for his help with his help in performing TFM on adhering cells.

## MATERIALS AND METHODS

### Cell culture

MRC-5, HeLa, and HEK 293T cells were cultured in Dulbecco’s modified Eagle’s medium (DMEM; D6429; Millipore Sigma, Darmstadt, Germany) supplemented with 10% fetal bovine serum (FBS; 12483020; Thermo Fisher Scientific, Waltham, MA, USA) and 0.1% Penicillin-Streptomycin (450-200-EL; Wisent, St-Bruno, QC, Canada). MDA-MB-231 and MDA-MB-468 cells were cultured in Leibovitz’s L-15 medium (41300-039; Thermo Fisher Scientific,Waltham, MA, USA) supplemented with 10% FBS without CO_2_ supplementation.

### Polysome profiling

Polysomal profiling was performed on MRC-5 cells at different time points during adhesion. Cells were washed with PBS, then lysed in 1 mL polysomal extraction buffer (20 mM Tris pH 7.5, 150 mM NaCl, 1.25 mM MgCl_2_, 5 U/mL RNase Inhibitor (M0314S; New England Biolabs, Ipswich, MA, USA), EDTA-free protease inhibitor cocktail, 1 mM DTT, and 1% Nonidet P-40) for 15 min on ice followed by 10 passes through a 10G needle.. Extracts were clarified by centrifugation at 12,000 × *g* for 20 min at 4°C. Cytoplasmic extracts were loaded on 15–55% (w/v) linear sucrose gradients for sedimentation by ultracentrifugation for 2,5 at 230,000 x g at 4°C. The gradient was read at 260 nm to generate a continuous absorbance profile. The area under the curve of the polysome fraction was calculated and compared to the area under the curve of the pooled 40S, 60S, and 80S fractions.

### Protein detection

Cells were extracted with Laemmli buffer (2% SDS, 2% β-mercaptoethanol, 20mM Tris pH 8.8, 10% Glycerol). For all cell lines, protein levels were quantified and normalized using the Bio-Rad RC DC^TM^ Protein Assay (5000122); Bio Rad, Hercules, CA, USA), and amido black staining was performed on each membrane to confirm protein transfer uniformity. Equal amounts of proteins were resolved by sodium dodecyl sulfate (SDS)-polyacrylamide gel electrophoresis (PAGE) and immunoblotted. Immunoblotting was performed with polyclonal antibodies against EIF2S1 (EIF2α; ab5369; Abcam, Cambridge, UK), EIF2S1 p-S51 (phospho eIF2α; ab32157; Abcam, Cambridge, UK), Talin (MAB1676; Millipore Sigma, Darmstadt, Germany), ⍺-Actinin (6487; Cell Signaling Technology, Danvers, MA, USA), Filamin-A (ab76289; Abcam, Cambridge, UK), Immunoreactive proteins were visualized with either goat anti-mouse (7076; Cell Signaling Technology, Danvers, MA, USA), goat anti-rabbit (7074; Cell Signaling Technology, Danvers, MA, USA) or TrueBlot anti-rabbit (Rockland Immunochemicals, Limerick, PA, USA) antibodies conjugated to horseradish peroxidase, and Western Lightning Plus Enhanced Chemiluminescence Substrate for Western Blotting (NEL104001EA; Perkin-Elmer, Waltham, MA, USA). Protein quantification was performed in ImageJ version 1.0 (US National Institutes of Health, Bethesda, MD, USA). GAPDH were used as negative controls.

### *In vitro* adhesion assays

Exponentially growing MRC-5 (*n*=3) and MDA-MB-231 (*n*=2) cells were detached using 0.025% trypsin with 0.1% ethylenediaminetetraacetic acid (EDTA), counted, and treated with either 1 µM ISRIB or 50 µg/mL cycloheximide. Control cells were left untreated. Samples were plated (HeLa: 15 000 cells/well, MRC-5: 25 000 cells/well, MDA-MB-468: 25 000 cells/well and MDA-MB-231: 50 000 cells/well) in 96-well Corning™ Stripwell™ Microplates (07-200-97; Fisher Scientific, Pittsburgh, PA, USA), and incubated at 37°C in 5% CO_2_ for 20, 40, 60, 90, 120, 240, and 360 min (*n* = 7 per time point). Cells were fixed with 3.7% formaldehyde for 15 min and nuclei were stained with DAPI (1 μg/mL). Fluorescence intensity was measured at excitation and emission wavelengths of 355 and 460 nm, respectively, using a Fluoroskan Ascent Microplate Fluorometer (5200110; Thermo Fisher Scientific, Waltham, MA, USA). Empty wells were used as controls for background subtraction. For the immunoblotting, exponentially growing MRC-5 (n=3), MDA-MB-231 (n=3) and MDA-MB-468 (n=3) cells were detached using 0,025% trypsin with 0,1% ethylenediaminetetraacetic (EDTA) resuspend in 12 mL of medium and agited during 20min at 37°C. After cells are seeded in 7 plates (one by time point) and mechanically lysated with Laemmli buffer at different times of adhesion (20,40,60,90,120,240 and 360 min) and one wash with 1mL of PBS at 20min was done to remove unadherent cells. For ISRIB (1μM), cycloheximide (50μg/mL) and Y27632 (10μM) we performed a pretreatment for 1 hour and the treatment was maintained all experiment long. For the contractility experiments, exponentially growing MDA-MB-231 and MDA-MB-468 cells were detached using 0,025% trypsin with 0,1% ethylenediaminetetraacetic (EDTA) resuspend in 12 mL of medium and agited during 20min at 37°C. After cells are counted and seeded in 7 or 9 flurodishs (one by time point and 70000 cells by fluorodish). One wash with 1mL of PBS at 20min was done to remove unadhered cells and cells were fixed with 3,7% formaldehyde for 10 min. For ISRIB (1μM), cycloheximide (50μg/mL) and Y27632 (10μM) we performed a pretreatment for 1 hour and the treatment was maintained all experiment long.

### Puromycin incorporation and immunoprecipitation assays

Exponentially growing cells were seeded in 100 mm plates (83.3902; Sarstedt, Newton, NC, USA) and incubated at 37°C in 5% CO_2_ for 45 min. Five minute prior to the time point completion, plates were washed carefully with prewarmed (37°C) complete medium to remove all unattached cells and incubated for the last five minutes with a prewarm complete medium (37°C) added with 5 μg/mL puromycin. Plates were washed with PBS and cells were scraped in 1 mL of lysis buffer (1% SDS, 5 mM EDTA, and 1 mM dithiothreitol (DTT)). After heating at 98°C for 5 min, the lysate was clarified by centrifugation at 12,000 × *g* for 10 min. The supernatant was diluted with 9 mL cold PBS and free puromycin was removed by centrifugation using Macrosep 3 kDa centrifugal filters (Pall). Concentrated lysate was diluted in non-denaturing lysis buffer (10 mM Tris pH 7.4, 1 nM EDTA, 150 mM NaCl, 1% Triton X-100, and EDTA-free protease inhibitor cocktail (Roche, 11873580001). Puromycylated peptides were immunoprecipitated using the 12D10 anti-puromycin antibody (MABE343; Millipore) bound to protein A agarose beads (11719408001, Millipore Sigma, Darmstadt, Germany), as previously done (Carbonneau et al., 2016, Huot et al., 2009). Samples were stored at-80°C until their analysis.

### MS data acquisition, analysis and archival

Peptide samples were analyzed at the Sherbrooke mass spectrometry core facility. Briefly, samples were loaded and separated using a Dionex Ultimate 3000 nanoHPLC system. 10 µL of the sample in 1% (v/v) formic acid was loaded with a constant flow of 4 µL/min onto a trap column (Acclaim PepMap100 C_18_ column, 0.3 mm id × 5 mm, Dionex Corporation, Sunnyvale, CA). Peptides were then eluted off to a PepMap C_18_ nano column (75 µm × 50 cm, Dionex Corporation, Sunnyvale, CA) with a linear gradient of 5–35% solvent B (90% acetonitrile with 0.1% formic acid) over a four hours gradient with a constant flow of 200 nL/min. Peptides were injected into an OrbiTrap QExactive mass spectrometer (Thermo Fischer Scientific, Waltham, MA, USA) using an EasySpray source. The voltage was set to 2.0 kV and the temperature of the column was set to 40°C. Acquisition of the full scan MS survey spectra (m/z 350–1600 was performed at a resolution of 70,000 using the accumulation of 1,000,000 ions. Peptides fragmented by collision-induced dissociation were selected on the ten highest intensities for the preview scan. The normalized collision energy used was 35%, and the resolution was set at 17,500 for 50,000 ions with a filling time set to a maximum of 250 ms for the full scans and 60 ms for the MS/MS scans. Unassigned charge states as well as singly, 7 and 8 charged species for the precursor ions were not analyzed, and a dynamic exclusion list was set to 500 entries over a retention time of 40 seconds under a 10 ppm mass window. The lock mass option was also enabled, and data acquisition was done using Xcalibur version 2.2 SP1.48. Protein identification and quantification were performed using the MaxQuant software package version 1.6.17.0 as described previously ^58^, with the protein database from UniProtKB (Homo sapiens, 29/01/2021, 77,027 entries). For protein identification, carbamidomethylation on cysteine was used as a fixed modification, and methionine oxidation and protein N-terminal acetylation were used as variable modifications. The enzyme was set to trypsin/P (*i.e.*, no cleavages on lysine or arginine before a proline), with up to two miscleavages allowed. The mass tolerance was 7 ppm for the precursor ions and 20 ppm for the fragment ions. All MS files were deposited at MassIVE (massive.ucsd.edu) and assigned the identified MSV000087439. They can be downloaded at ftp://MSV000087439@massive.ucsd.edu.

### Quantitative polarization microscopy

The polarization microscope is built on an inverted Axiovert microscope equipped with a Zeiss Axiocam 305 camera. A linear polarizer actuated by a motorized rotation stage (Thorlabs) was positioned in the illumination plane above the condenser, and a circular polarizer (analyzer) was positioned in the imaging plane. Images were acquired using a 20x 0,9 NA polarization objective (Zeiss). Image sequences were acquired with 5° intervals of the rotating polarizer over a range of 0 to 180 degrees using Zen software. Acquisition time for each individual picture was set at 20ms. A timed delay of 2 seconds was used between each acquisition to allow rotation of the polarizing element. The polarized image sequences were then processed with a custom Python code to obtain a pixel-by-pixel retardance map. The resulting retardance images were then background subtracted and analyzed with ImageJ. The retardance of the whole cell was quantified as the average value of the background-subtracted retardance of cell area. Generated data points are from at least 3 independent experiments for both MDA-MB-231 (n=50 cells) and MDA-MB-468 (n=50 cells).

### Traction force microscopy

The traction force microscopy was performed as described previously (PMID: 38194965). MDA-MB-231 cells were detached using TrypLE (Gibco, 12605010) and resuspended at 50 000 cells/mL in DMEM (Corning, MT10013CV), 10% FBS (Gibco, A5256701), Antibiotic-Antimycotic (Gibco, 15240062) and 1ug/ml WGA 488 (Invitrogen, W6748). Cells were then rotated for 20 min at 12RPM at 37oC. A polyacrylamide hydrogel with a stiffness of 16 kPa containing 0.04 um fluorosphere (Thermofisher, F8789) was placed in a custom magnetic cell chamber with 1x PBS (Corning, 21,040 CV) covering the hydrogel to prevent it from drying. The empty cell chamber was then installed on a Spinning Disk (W1 Confocal Spinning Disk, Yokogawa) to adjust the focal plan to the top of the gel. Once the set-up is finished, the PBS was removed and 1 ml of cell suspension was added to the gel. Twenty fields were chosen; then cells and gels were imaged every 2 minutes post-seeding for 6 hours at 37oC with 5% Co2. Cells were then removed with 0.05% SDS (RPI, L22010-500) to take the reference image of the gel. TFM analysis was performed based on the protocol described by Schmitt et al. 2024 ^59^. For an accurate measurement of forces, only isolated cells were kept for analysis.

### Computational modelization of mechanical stress during cell adhesion

*Tensegrity model:* To establish the force and stress distribution within the cell following its adhesion, a computational model based on tensegrity was developed in Matlab (Mathworks). Basis for cellular tensegrity are already described ^60,61^. Briefly, the cell can be modeled as compressive struts (microtubules) that counterbalance tensional stress from prestressed cables (actin filaments and intermediate filaments). Based on the organization and interconnection within the system, a linear equation system can be established to describe the stresses density at every node:

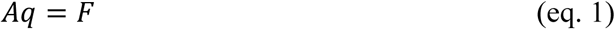

Where A is a equilibrium matrix, q is the internal forces density matrix and F is the external applied force. The equilibrium matrix ca further be defined by a connectivity matrix linking each node with its associated structural bars:

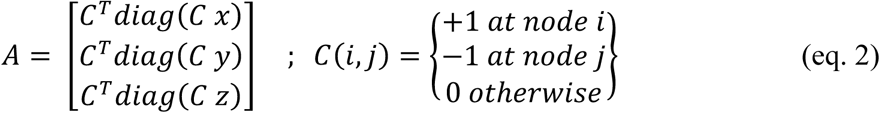

The equilibrium matrix is related to the generalized stress σ when it is under a generalized load f:

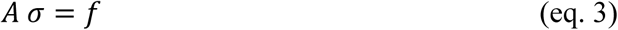

This equation system can then be used to determine if the structural components are stable and extract the internal stress information and the total force exerted by all the components. We assume an initial icosahedron configuration for the cells with 12 nodes and 24 rods. To alter the structure, new nodes were generated at every timepoints following a logistic distribution to match cell spreading and focal adhesion formation. An equivalent logistic distribution was used to model the probability of a node being pruned (simulating an adhesion disassembly). Moreover, each actin microfilament cables were allowed to be crosslinked with actin bundling proteins (e.g. Filamin or ⍺-Actinin) or converted to a stress fiber, effectively increasing the stiffness and bundling rods assigned as actin components. The stiffening resulting from increased actin crosslinking and activation of the actomyosin mediated force was modeled following a logistic regression whose maximal parameters were determined according to known stiffness of actin bundles or stress fiber structures. The corresponding logistic regression at time t and scaled to the initial and final stiffness is given by:

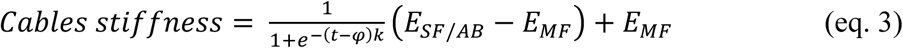

Where φ is the time delay to reach 50% crosslinking, k is an estimated crosslinking and fiber bundling rate, E_MF_ E_AB_ and E_SF_ are the effective modulus for actin microfilaments (i.e. the assembled actin scaffolds, not an individual fiber), crosslinked actin bundles and stress fibers respectively. Of note, the k value for stress fibers was set lower to account for the slow mechanism of bundling together numerous actin cables. The traction force |T| and shear stress τ within each rod structure was computed at every time step from the strain-stress relationships:

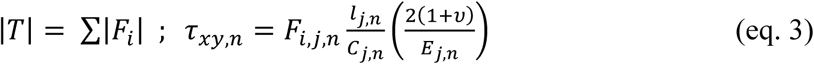

Where, *F*_i,j,n_ is the force acting on node *i*; *l*_j,n_, *C*_j,n_, *E*_j,n_ and ν are respectively the length, crossectional area, Young’s modulus and Poisson’s ratio of the cytoskeletal component connected to node *i*. The parameters for the tensegrity model are provided in table 1. When possible, the parameters were estimated from the literature. In some case (K_myosin_), the activation rate was set lower than available values since RhoA activation is known to be inhibited during early adhesion, peaking between 60 and 90 min ^62^.

**Table 1:** Tensegrity modelization parameters.

| Variable |  | Value | Units | REF |
| --- | --- | --- | --- | --- |
| $\alpha_{\text{node}}; \beta_{\text{node}}$ | Gamma distribution parameters for node formation during cell spreading | 1; 50 | NA | Est. |
| $\alpha_{\text{prune}}; \beta_{\text{prune}}$ | Gamma distribution parameters for node pruning during cell spreading | 7; 15 | NA | Est. |
| $K_{+\text{node}}$ | Cytoskeletal nodes creation at steady state | 0.05 | $\text{min}^{-1}$ | Est. |
| $K_{\text{prune}}$ | Cytoskeletal nodes pruning at steady state | 0.033 | $\text{min}^{-1}$ | Est. |
| $Df_0$ | Initial actin stress density at $t = 0$ | 0.3 | $\text{nN}/\mu\text{m}$ | Est. |
| $Df_{\text{max}}$ | Maximal actin force density | 0.4 | $\text{nN} \cdot \mu\text{m}^{-1}$ | Est. |
| $\phi_{AB}, \varphi_{AB}$ | Logistic central parameters | 80 | min | Est |
| $K_{AB}$ | Actin bundling/crosslinking rate | 0.0667 | $\text{min}^{-1}$ | 63,64 |
| $K_{SF}$ | Stress fiber crosslinking rate | 0.0167 | $\text{min}^{-1}$ | 65 |
| $K_{\text{myosin}}$ | Signaling induced Myosin activation rate | 0.05 | $\text{min}^{-1}$ | 66 |
| $E_{MF}$ | Microfilament Effective Elastic Modulus | $10^2$ | Pa | 67-69 |
| $E_{AB}$ | Actin bundle Effective Elastic Modulus | $10^4$ | Pa | 69-71 |
| $E_{SF}$ | Stress fiber Effective Elastic Modulus | $10^6$ | Pa | 72,73 |
| $E_{IF}$ | Intermediate filament Effective Elastic Modulus | $10^{10}$ | Pa | 74 |
| $E_{MT}$ | Microtubule Effective Elastic Modulus | $10^9$ | Pa | 75 |
| $A_{MF}$ | Microfilament cross section | 0.0001 | $\mu\text{m}^2$ | 76 |
| $C_{SF}$ | Stress fiber cross section | 0.015 | $\mu\text{m}^2$ | 72,73 |
| $C_{IF}$ | Intermediate filament cross section | 0.0002 | $\mu\text{m}^2$ | 76 |
| $C_{MT}$ | Microtubule cross section | 0.0005 | $\mu\text{m}^2$ | 76 |

*Clutch model:* To modelize Talin dynamic and breakage in focal adhesions, a modified clutch model based on a Matlab code publicly available was used ^28^. Briefly, a script was used to run the motor clutch computational simulation based on the output of the tensegrity computational model, including the stress and a number of focal adhesions (here approximated as the number of nodes in contact with the surface). The action of eif2α on protein production was taken into account by modeling a 2 phases response. First, a logistic decay was assumed for eif2 phosphorylation to go back down to steady state levels after 2 hours. Second, a gaussian fit was assumed to match early peak activation. The parameters of each fit were adjusted based on the experimental data. The phosphorylation signal from eif2 was then used to proportionally block Talin production rate. The parameters for the clutch model were based on the original code as listed in Table 2. The clutch model was run for 20 iterations for each adhesion (each iteration corresponding to 1 sec). In addition, the state of Talin binding to both actin and integrin (clutch on) was recorded and passed on to subsequent time point. A peak force of 100 nN acting on a single Talin molecule was considered as the threshold to induce irreversible damage on the protein. The number of damaged Talin proteins for each call of the clutch model was provided as the main output. Moreover, since protein production is a tightly regulated process to maintain homeostasis, a PID was implemented to prevent an overproduction of Talin proteins.

**Table 2.**
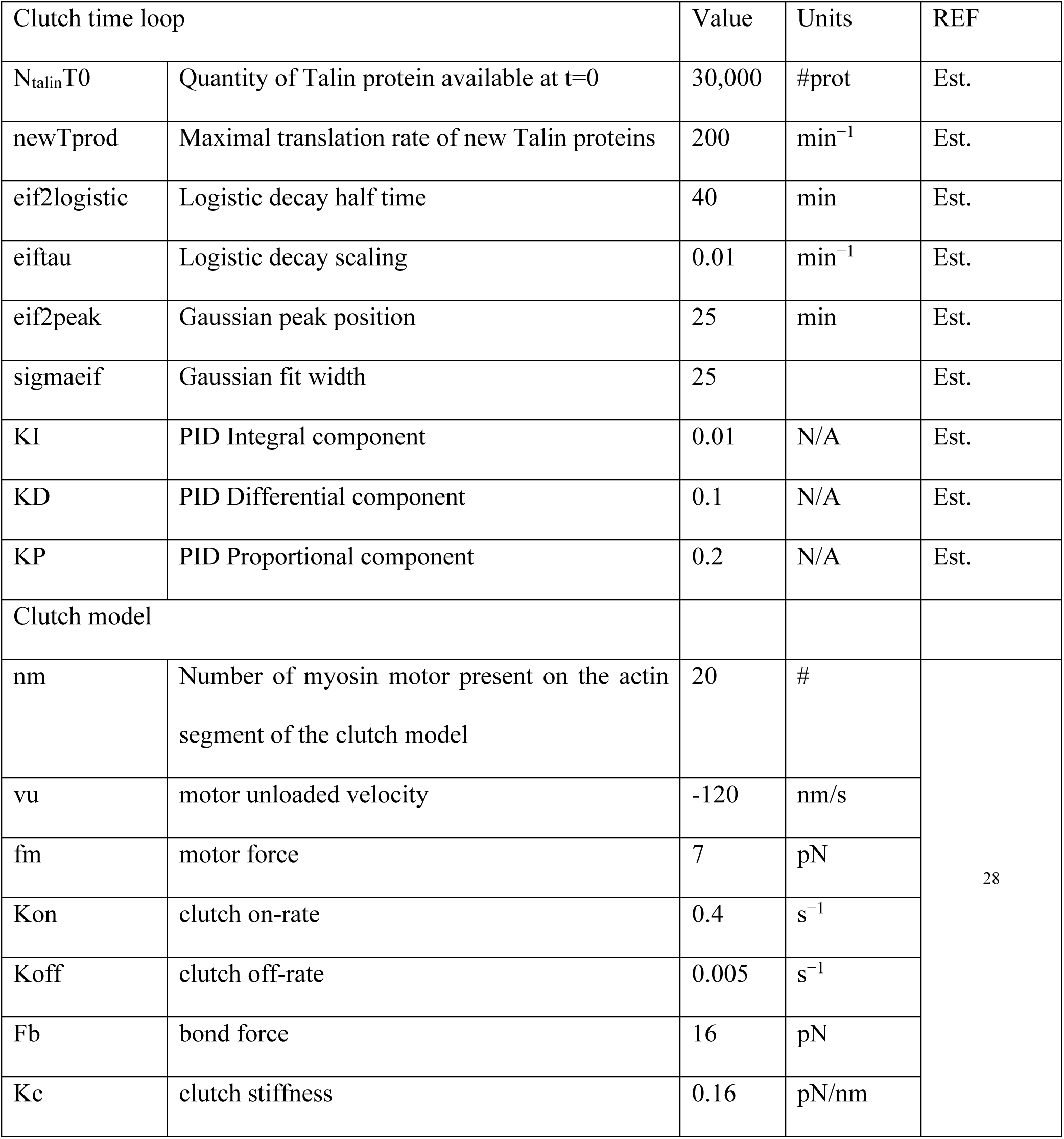
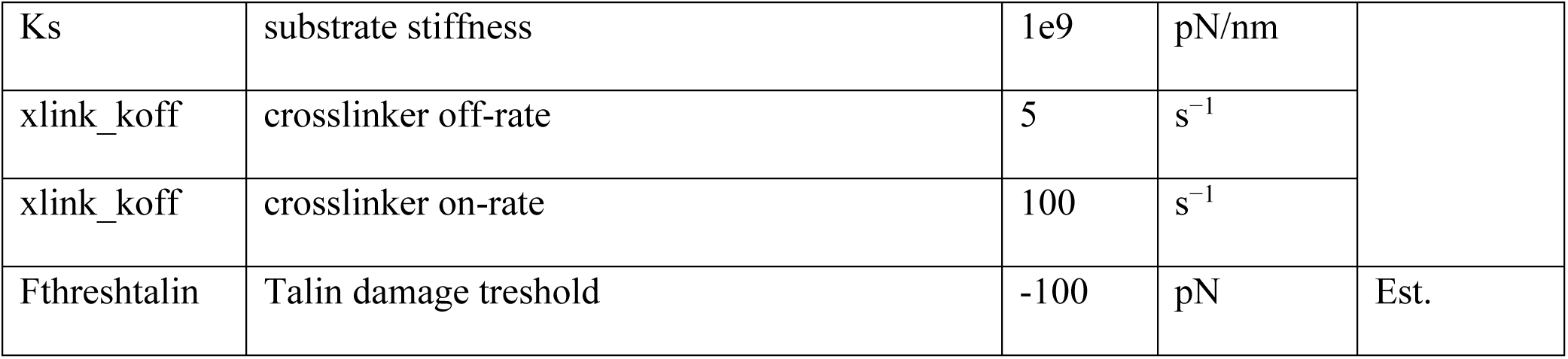

| Clutch time loop |  | Value | Units | REF |
| --- | --- | --- | --- | --- |
| $N_{\text{talinT0}}$ | Quantity of Talin protein available at $t=0$ | 30,000 | #prot | Est. |
| newTprod | Maximal translation rate of new Talin proteins | 200 | $\text{min}^{-1}$ | Est. |
| eif2logistic | Logistic decay half time | 40 | min | Est. |
| eiftau | Logistic decay scaling | 0.01 | $\text{min}^{-1}$ | Est. |
| eif2peak | Gaussian peak position | 25 | min | Est. |
| sigmaeif | Gaussian fit width | 25 |  | Est. |
| KI | PID Integral component | 0.01 | N/A | Est. |
| KD | PID Differential component | 0.1 | N/A | Est. |
| KP | PID Proportional component | 0.2 | N/A | Est. |
| Clutch model |  |  |  |  |
| nm | Number of myosin motor present on the actin segment of the clutch model | 20 | # | 28 |
| vu | motor unloaded velocity | -120 | nm/s |  |
| fm | motor force | 7 | pN |  |
| Kon | clutch on-rate | 0.4 | $\text{s}^{-1}$ | |
| Koff | clutch off-rate | 0.005 | $\text{s}^{-1}$ | |
| Fb | bond force | 16 | pN |  |
| Kc | clutch stiffness | 0.16 | pN/nm |  |
| Ks | substrate stiffness | 1e9 | pN/nm |  |
| xlink_koff | crosslinker off-rate | 5 | s<sup>-1</sup> |  |
| xlink_koff | crosslinker on-rate | 100 | s<sup>-1</sup> |  |
| Fthreshtalin | Talin damage treshold | -100 | pN | Est. |

## Code availability

The tensegrity and modified clutch Matlab code (R2021a, Mathworks) used to make the computational models are available from the corresponding author (FB) upon reasonable request. The original clutch model is available at: https://oddelab.umn.edu/software. The TFM code used in this study can be found at: https://github.com/OakesLab/TFM.

---

**Fig. S1.**
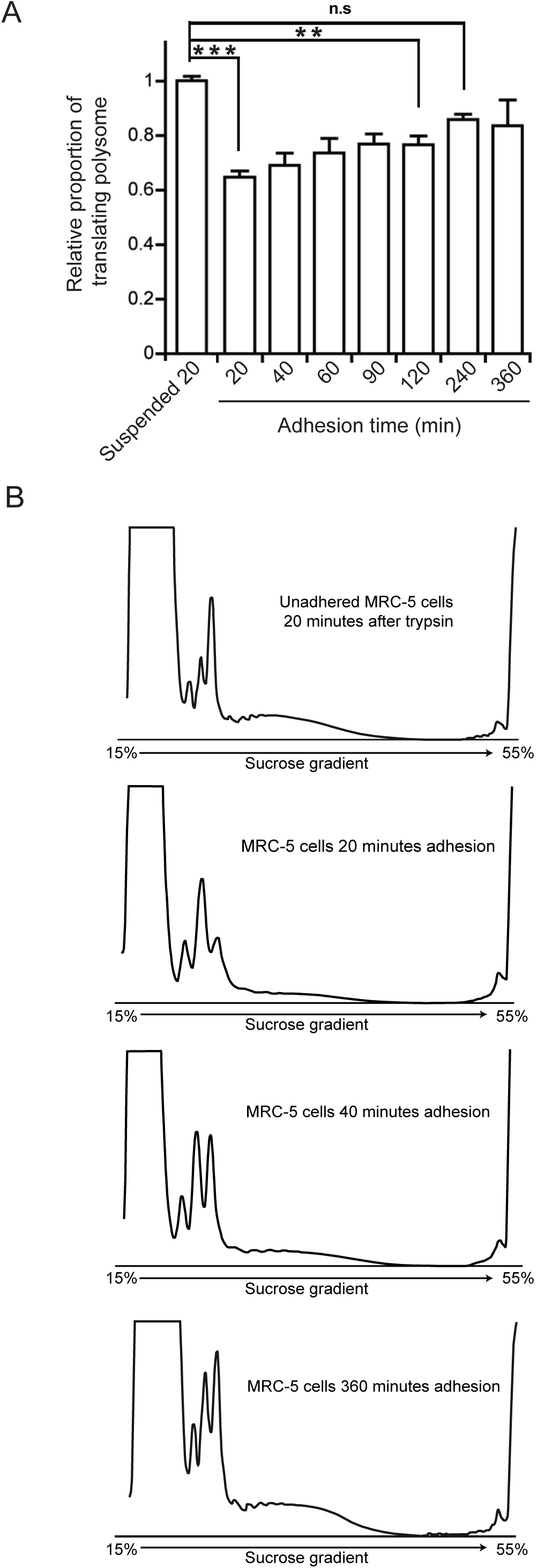
Relative translation activity throughout the adhesion process **(A)** Quantification of the heavy polysomal fraction corresponding to translating polysome in MRC-5 cells at different time points throughout the adhesion process. Extracts were loaded on 15–55% (w/v) linear sucrose gradients and the relative proportion of translating polysome was measured using the area under the curve of the polysomal fractions over the area under the curve of the pooled 40S, 60S, and 80S fractions. **(B)** Polysomal profile of resuspended cells maintained in suspension for 20 minutes and cells that were adhering for 20, 40 and 360 minutes. The latter time points represent cells that are fully adhered.

**Fig. S2.**
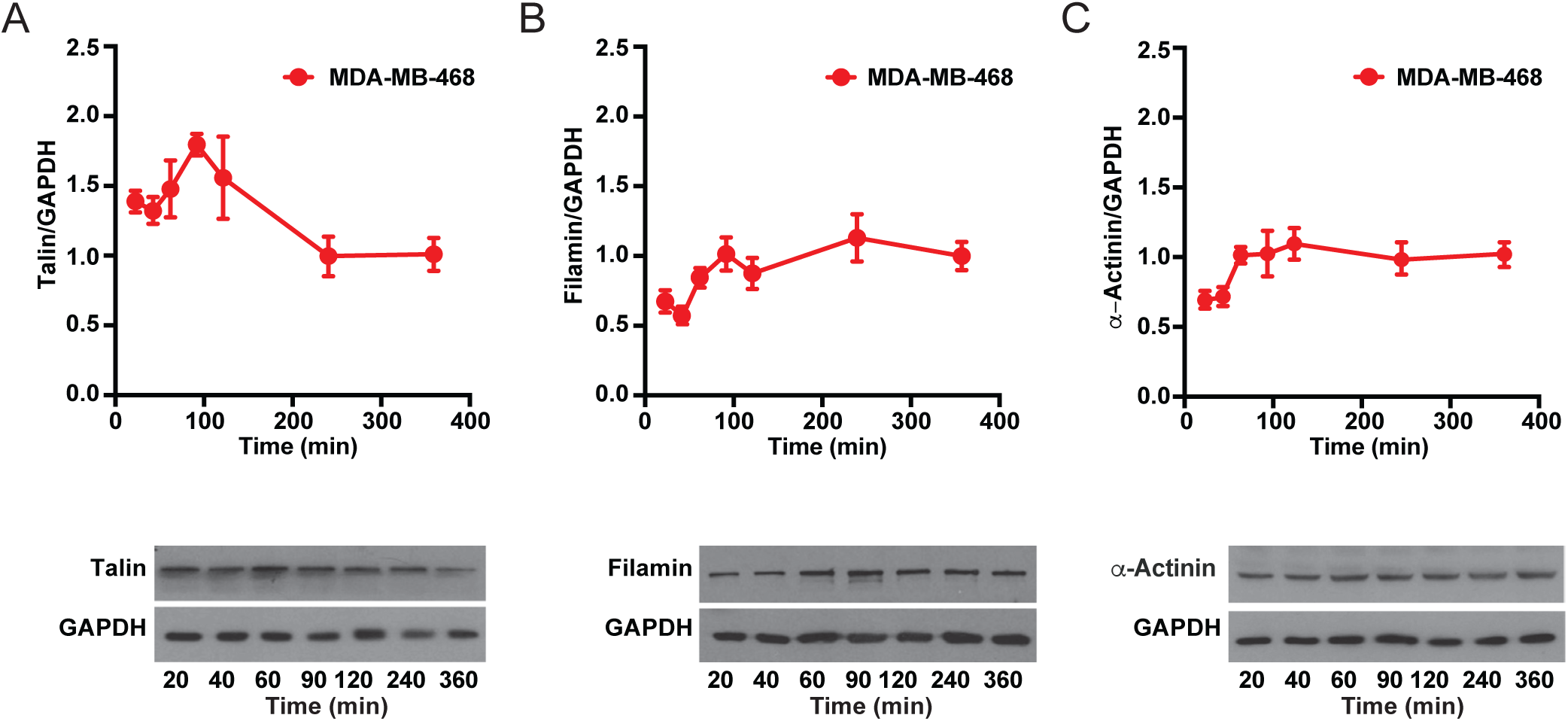
Expression of adhesion-and cytoskeleton-related proteins during MDA-MB-468 adhesion. Quantification of levels of translated adhesion related mRNAs of Talin (**A**), Filamin-A (**B**), and ⍺-Actinin (**C**), at different time points during cellular adhesion in MDA-MB-468 untreated. Lower panels show representative western blot detection of each protein at different time points during cellular adhesion in in MDA-MB-468 cells. Protein expression was normalized at time point 360 minutes, as we found that adhered cells were morphologically mature independently of treatments.

**Fig. S3.**
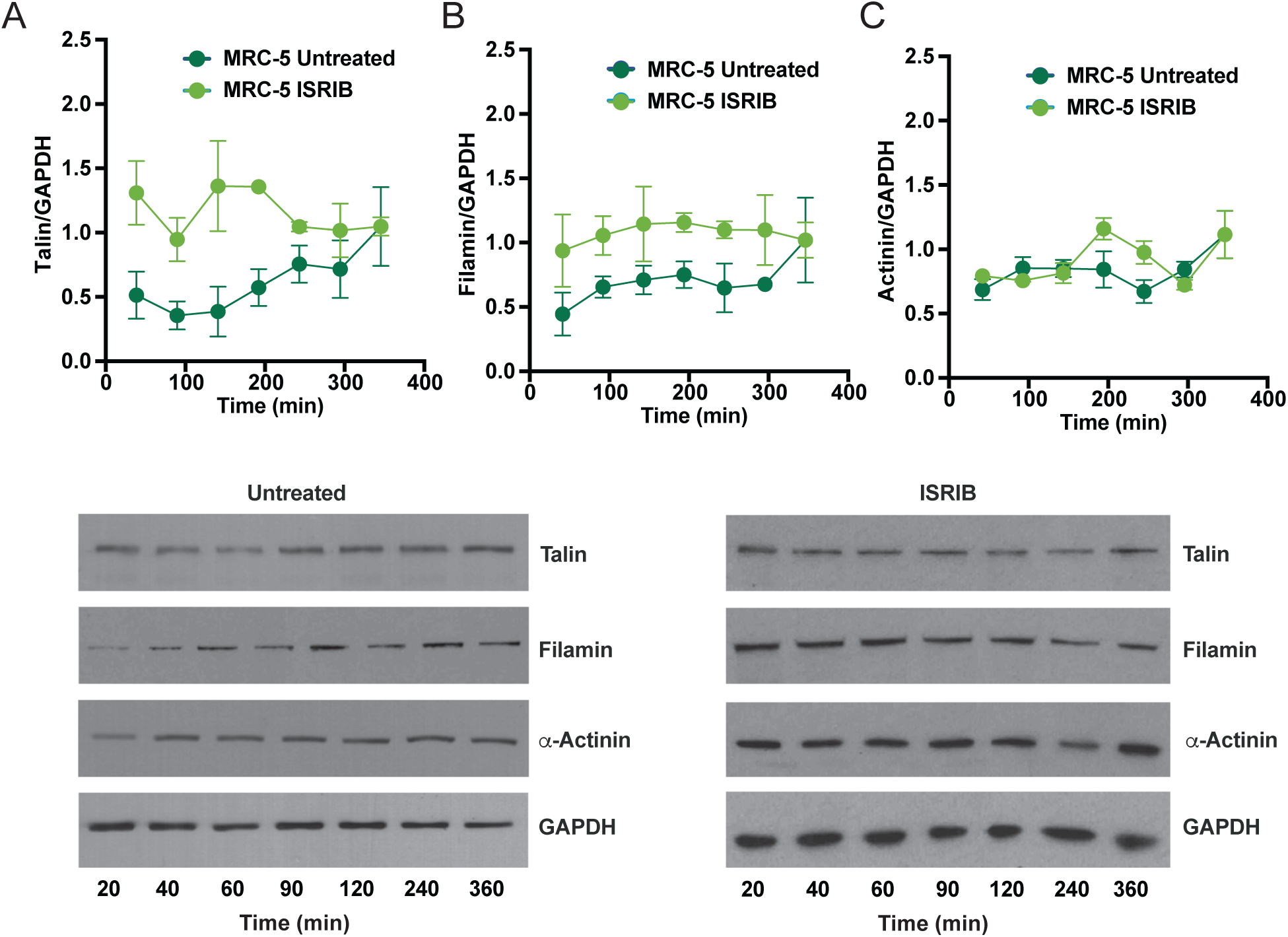
Expression of adhesion-and cytoskeleton-related proteins during MRC-5 adhesion. Quantification of levels of translated adhesion related mRNAs of Talin (**A**), Filamin-A (**B**), and α**-**Actinin (**C**), at different time points during cellular adhesion in untreated MRC-5 cells (DMSO) or treated with 1μM ISRIB (n=3). Lower panels show representative western blot detection of each protein at different time points during cellular adhesion in untreated MRC-5 cells (DMSO) or ISRIB-treated MRC-5 cells (1μM). Protein expression was normalized at time point 360 minutes, as we found that adhered cells were morphologically mature independently of treatments.

**Fig. S4.**
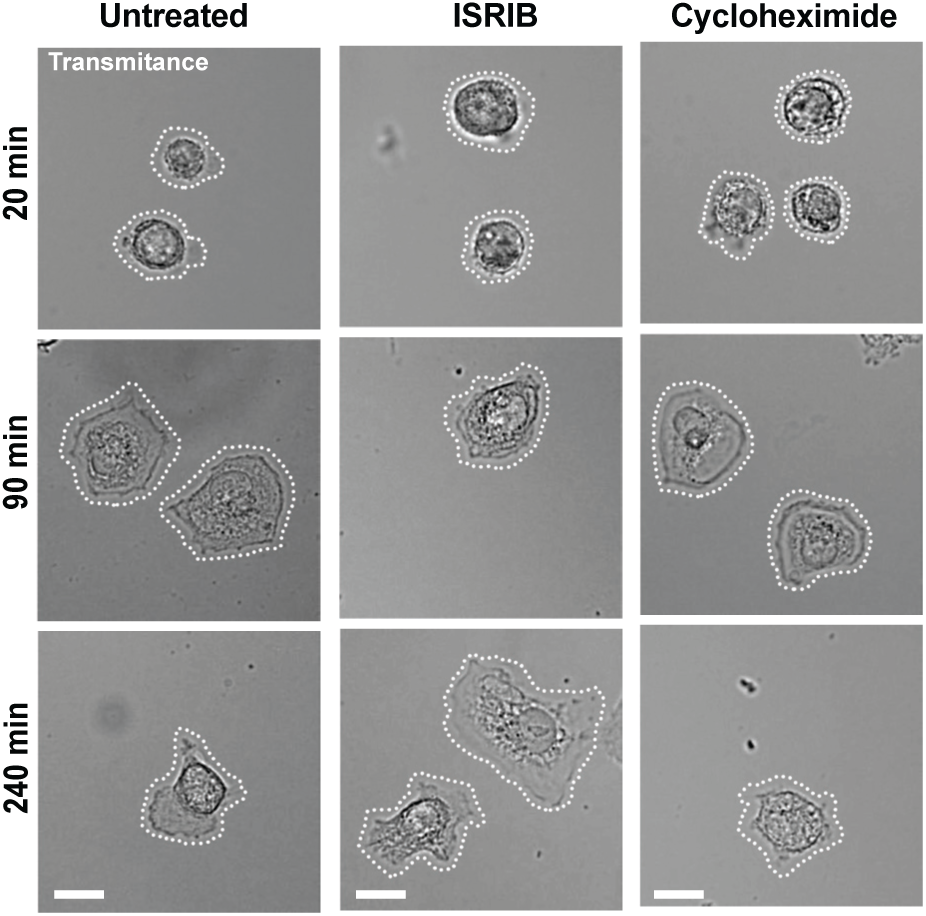
Transmitted light images of MDA-MB-231 cells Representatives transmitted light images of untreated MDA-MB-231 cells (DMSO) or MDA-MB-231 treated with either cycloheximide (50μg/mL) or ISRIB (1μM) at 20 min, 90 min and 240 min post-seeding. Scale bar = 20 μm.

